# Investigating TRSV Phytoalexin Infusions as a Novel Treatment for Alzheimer’s Disease Using ApoE4 Plasmid Transfections in Neuro2a Cell Complexes

**DOI:** 10.1101/2020.04.05.026674

**Authors:** Om Gandhi

## Abstract

Because the accumulation of amyloid plaques plays a central role in the pathogenesis of Alzheimer’s disease (AD) and is responsible for many of its neurodegenerative effects, this project aims to evaluate the effectiveness of TRSV phytoalexin infusions as a novel treatment for the disease by comparing three separate agents and determining its impact on the accumulation of amyloid plaques in mouse Neuro2a cells. The experimental model is as follows: Neuro2a cells were first thawed using incubation and centrifugation techniques. The subculturing protocol was then performed to maximize cellular viability using DPBS and TrypLE dissociation reagents. The cells were transfected with pCMV4-ApoE4 bacterial plasmid using Opti-MEM medium and Lipofectamine LTX while also being treated with phytoalexin agents. Bradford’s assay was performed on the samples using CBB G-250 reagent, and they were run through a spectrophotometer and a Python low-pass filter to generate readings. The experimental data showed that natural grape seed phytoalexin extract was the most effective treating agent, followed by curcumin extract and synthetic RDS phytoalexin. All treating agents lessened the accumulation of the amyloid plaques significantly, supporting the novel infusion protocol as a treatment for AD.

## 1. Introduction

Alzheimer’s disease (AD) is a neurodegenerative condition that results in the continuous decline of one’s cognitive, behavioral, and social capabilities [1]. It is the most common type of dementia that slowly destroys memory along with the ability to carry out both simple and complex tasks [1,2]. In total, AD currently afflicts about an estimated 5.8 million Americans age 65 and older and is the sixth leading cause of death in the United States. It is also the leading cause of mental degeneration in the elderly and accounts for a large percentage of admissions to long-term care facilities [3]. Despite there being no way to stop the progression of the disease completely, advancements in scientific research have allowed for the understanding of AD at the cellular and molecular level.

### 1.1. The Biological Basis of AD

The healthy human brain consists primarily of tens of billions of neurons that process and transmit information via electrical and chemical signals, regulating everything from muscle movement to energy storage. At first, AD disrupts connections between neurons involved with memory in areas such as the hippocampus, resulting in loss of function and cellular death [4]. It later affects areas in the cerebral cortex responsible for language, reasoning, and social behavior. Eventually, many other regions of the brain are damaged, leading to death [4,5].

What can explain the brain damage commonly associated with AD? Although the definitive causes and mechanisms of the disease have yet to be determined, recent studies point to a series of common factors. A range of research has shown that the neurodegenerative phenomena are initiated and enhanced by oxidative stress, a state characterized by an excessively high ratio of oxidants to antioxidants [6–10]. This imbalance can occur either as a result of decreased oxidation defense or an increase in the concentration of free radicals, commonly defined as compounds with unpaired electrons in their valence shells. Free radicals are well known as highly-reactive health hazards, but they attack some types of molecules more than others [11].

The impact of oxidative stress does not stop with the brain’s physical structure. High concentrations of reactive oxygen species (ROS), such as free radicals, also degrade enzymes and proteins that are crucial to normal neuronal and glial cell function [12]. This is the case for two enzymes, which are especially sensitive to oxidative modification: glutamine synthetase and creatine kinase, both of which are less present in AD brains. Oxidative impairment of creatine kinase may also cause decreased energy metabolism, demonstrating the critical nature of the enzymes that AD-related oxidative stress hurts most [12,13]. AD also results in the abnormal deposition of neurofibrillary tangles, “characterized by the aggregation and hyperphosphorylation of the *τ* protein into paired helical filaments,”[14]. The pathologic aggregation of the *τ* protein leads to fibril formation, insolubility, and an inability for affected neurons to send signals to other neurons, leading to neurological degeneration [14,15]. Furthermore, phosphorylation is linked to oxidation through the microtubule-associated protein kinase pathway and activation of the transcription factor nuclear factor-*κ*B (NF-*κ*B), thus linking oxidation to the hyperphosphorylation of *τ* proteins [14–17]. Perhaps the most significant biological indicator of AD, however, is the presence of amyloid plaques.

The formation of amyloid plaques requires two specific proteins: beta-amyloid protein and apolipoprotein E (APOE) [18]. While the beta-amyloid protein is the primary protein involved, APOE promotes the accumulation of beta-amyloid proteins into plaques. These insoluble plaques have toxic properties, including synaptic dysfunction, *τ* phosphorylation, and neurodegeneration, that contribute significantly to the progression of AD [18,19].

### 1.2. Beta-Amyloid Protein

The beta-amyloid (A*β*) 4-*κ*Da peptide has a length that varies from 38 to 43 amino acids. The protein itself usually is present in a variety of somatic cells, but its normal biological function remains unknown [20]. “Under physiological conditions, A*β*40 constitutes about 90 percent of the total amount of A*β*,” [20]. Of the two significant species of A*β* (A*β*40 and A*β*42), the latter is more aggregation-prone due to two additional hydrophobic amino acids [20,21]. This accounts for A*β*42’s position as the predominant compound accumulating in plaques in the AD brain.

Abnormal deposition of the A*β* protein is caused by aggregation of the transmembrane amyloid precursor protein (APP), which is synthesized by the sequential action of *β*-secretase (BACE1) and *γ*-secretase [22–25]. “Amyloid precursor protein (APP) is an integral membrane protein, with a large N-terminal extracellular domain and a short C-terminal cytoplasmic domain,” [22]. It is synthesized in the endoplasmic reticulum (ER), post-transcriptionally modified in the Golgi apparatus, and then transported to the cell surface, where it is attached to the plasma membrane. APP is expressed throughout the body, and A*β* is a typical product of APP metabolism. “The APP gene is spliced to yield isoforms of various lengths; the 751- and 770-isoforms predominate in non-brain or nerve cells, whereas the 695-amino acid form (APP695) is by far the most predominant form in neurons,” [22]. APP and APP-like protein have orthologues across nearly all animals, indicating that they may be necessary for a variety of biological processes. However, their exact functions are not currently known. “Possible roles for APP and its proteolytic products range from axonal transport to transcriptional control and from cell adhesion to apoptosis,” [24].

Though comparatively little may be known about the cellular function of APP, much is known about its genetics and proteolytic processing, particularly concerning AD. APP is first cleaved by either *α*- or *β*-secretase at the respective *α*- or *β*-sites [20,25–27]. These two sites are near the extracellular side of APP’s transmembrane domain, allowing for more efficient cleavage. *α*- and *β*-secretase proteases compete for APP cleavage to give two products: a soluble APP and a membrane-anchored C-terminal stub [20,25–29]. This stub is the site of the crucial AD indicated cleavage, where *γ*-secretase acts. “While the division of C83 generates the 6-*κ*Da APP intracellular domain (AICD) and releases the N-terminal 3*κ*Da peptide (p3) into the extracellular space, cleavage of C99 generates AICD and the A*β* peptide,” [20]. The processing of APP by beta-secretase and gamma-secretase and the corresponding formation of beta-amyloid protein is known as the amyloidogenic pathway. In the alternative path of APP processing termed as the non-amyloidogenic pathway, no A*β* is produced as *α*-secretase cleaves APP with an A*β* sequence [27–30]. The second type of protein required for the formation of amyloid plaques is the Apolipoprotein E.

### 1.3. Apolipoprotein E (APOE)

The ApoE gene has demonstrated to code for Apolipoprotein E, a fat-binding protein. Although everyone inherits a copy of some form of the APOE gene from each parent, it is estimated that between 40 and 65 percent who are diagnosed with AD have the ApoE4 gene, implying a genetic basis for the disease [31]. There are three common forms of the APOE gene: ApoE2, ApoE3, and ApoE4. “ApoE has an N-terminal domain and a C-terminal domain, which are the receptor-binding region and fat-binding region, respectively,” [32]. There are three different isoforms, APOE2, APOE3, and APOE4. The structural differences between them are located at amino acid residues 112 and 158. While ApoE2 has cysteine 112 and cysteine 158, ApoE3 has cysteine 112 and arginine 158, and ApoE4 has arginine 112 and arginine 158. The changes in the R-group of amino acids result in structural and functional (i.e., binding ability) differences between them [32,33].

The exact mechanism connecting APOE isoforms and A*β* metabolism is not yet clear. Recent studies, with evidence collected from both neuron cultures and mice models, have identified that ApoE could activate the DLK-MKK7-ERK1/2 cascade, which stimulates transcription factor AP-1 [33–35]. Increased activity of AP-1 eventually results in a higher rate of APP transcription and, as previously discussed, elevated A*β* production. ApoE4 was shown to have the highest activation of this cascade and, in turn, generated more A*β* than APOE3 and APOE2. A*β* also exhibited accelerated fibrillization when incubated with ApoE4, when compared to ApoE3 in vitro [35,36]. Moreover, extracellular ApoE positively affected the targeting of lipid rafts with A*β*, where it was converted to an oligomer. As previously mentioned, the brain is primarily composed of vulnerable lipid-based substances. Extracellular ApoE can direct A*β* protein to bind to fatty substances, like those found in the brain [37]. Therefore, ApoE4 can assist in the formation of A*β*, playing a crucial role in the accumulation of beta-amyloid proteins that results in the nefarious amyloid plaques. Since these plaques play such a central role in the pathogenesis of AD [18,19], it is vital to investigate how to limit their accumulation.

### 1.4. Trans-Resveratrol

Resveratrol is a natural phytoalexin that acts against pathogens and is an integral part of the immune system in more than 70 plant species [38]. Its unique structure as a 3,5,4′-trihydroxy-trans-stilbene possessing two phenyl rings linked to each other by an ethylene bridge allows it to possess a variety of properties and capabilities [38,39].

“As a natural food ingredient, numerous studies have demonstrated that resveratrol possesses a very high antioxidant potential,” [38]. This makes it a prime candidate for the restoration of the correct balance between oxidants and antioxidants, thus limiting oxidative stress, one of the neurodegenerative factors present in AD. Resveratrol also exhibits antitumor activity, and its anti-cancer properties contribute significantly to its ability to inhibit all carcinogenesis stages [38–40]. Other positive bioactive effects have also been reported. These effects stretch from anti-inflammatory action to cardioprotection, and from vasorelaxation to neuroprotection. Although resveratrol and trans-resveratrol are similar in their many characteristics, one defining feature of trans-resveratrol is its ability to act as a free radical scavenger, stabilizing harmful unpaired valence electrons and decreasing damage to body tissues [40]. This makes the compound especially promising as a potential treatment agent for AD.

The primary purpose of the project is to determine whether grape seed phytoalexin extract or curcumin phytoalexin extract serves as the most effective natural trans-resveratrol (TRSV) treating agent by best limiting the accumulation of amyloid plaques. This project also aims to compare the treatment capabilities of the natural TRSV phytoalexins to a chemical phytoalexin supplement: resveratrol dietary supplement (RDS).

## 2. Results and Discussion

### 2.1. Viability Rates

The viability rate calculation determines if the living cultured cells are evenly distributed throughout the flask, a crucial step in minimizing errors as well as creating a model representative of human neurophysiology to maximize practical applications. Figure 1 shows the viability rates of the subcultured Neuro2a cells before any plasmid transfection or phytoalexin treatment.

**Figure 1.**
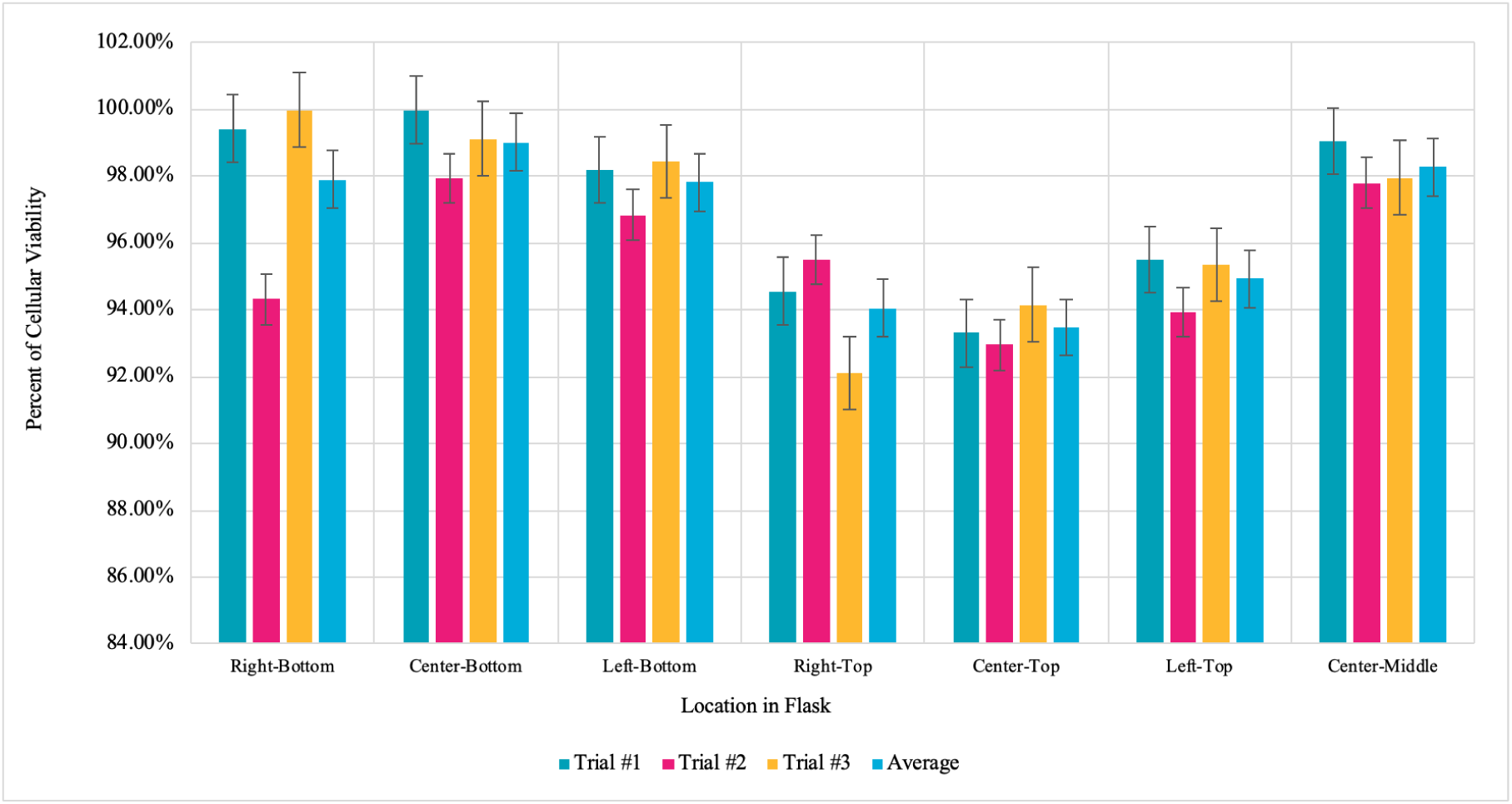
Neuro2a cell viability rates at different locations in the flask: right-bottom, center-bottom, left-bottom, right-top, center-top, left-top, center-middle. The viability was measured three times in each location to serve as trials and to determine consistency. Error bars were set at standard error.

Since all of the viability rates are higher than 90 percent, as seen in Figure 1, the thawing and subculturing methods were successful in maximizing the vitality of the cells. However, there is some difference between the viability rates of the cells at different locations in the flask. While the cells at the bottom had an average viability rate of around 98.24 percent, the cells at the top of the flask had a viability rate of 94.15 percent. This is primarily due to the top cells having more exposure to contaminants that may hinder their biological processes, such as metabolism or reproduction. Nonetheless, the cells used in the experimentation were taken from the bottom of the flask to maintain a realistic model.

### 2.2. Amyloid Concentrations in Cells with Phytoalexin Treatment

The concentration of amyloid plaques in the cells infused with the different sources of the natural phytoalexin extract was measured over a time interval of 400 seconds. Using Python, the average of the values for a specific trial over the time interval was calculated. The average of all the data points across all trials was also calculated.

According to the experimental data of Figure 2a, the average concentration of amyloid plaques across all trials was 8.623 pg/mL when the cells were infused with curcumin extract. Despite there being some deviation between the trials, the standard deviation and standard error of the samples are relatively low, especially considering that the unit of measurement is picograms. With an average of 5.444 pg/mL across all trials in Figure 2b, the cells infused with the grape seed phytoalexin extract had the lowest concentration of amyloid plaques measured in this experiment. However, there is some deviation in the data, especially in the graph. This can be attributed to uncontrollable errors, such as microscopic contaminants on the sides of a cuvette. Nonetheless, the data is relevant due to a low calculated standard deviation and standard error.

**Figure 2.**
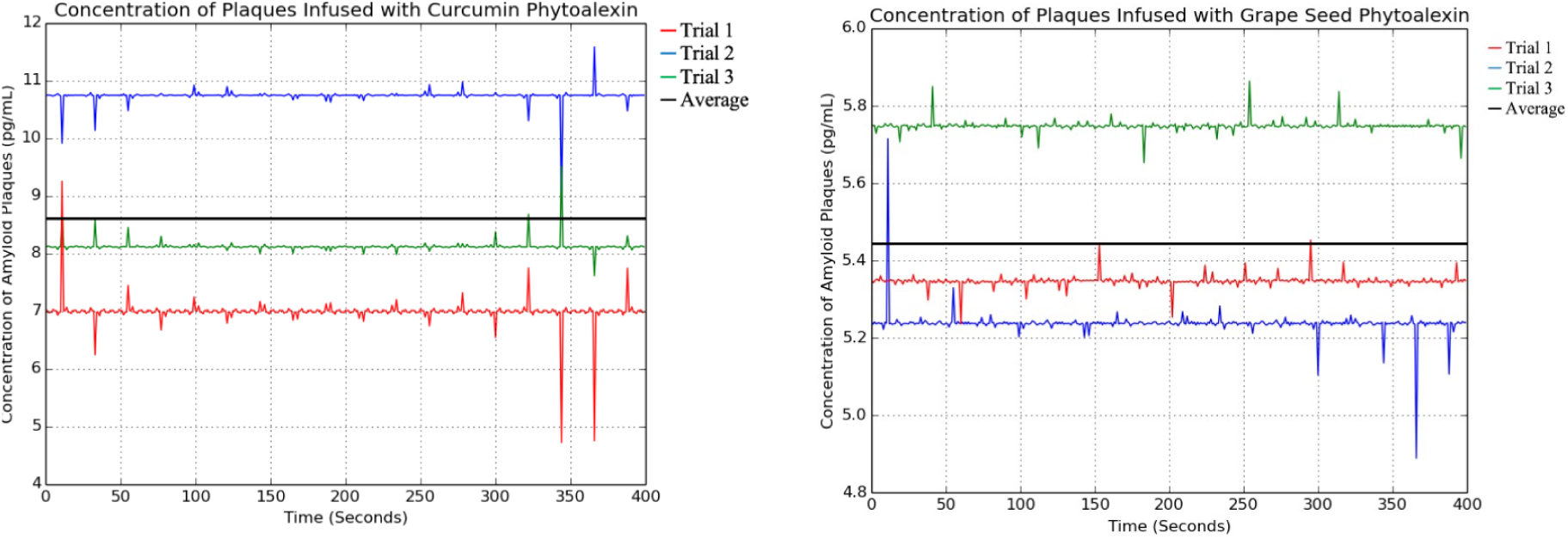
The concentration of amyloid plaques in the cells infused with the curcumin phytoalexin extract (**a**) and the grape seed phytoalexin extract (**b**) extract over a time interval of 400 seconds.

Per Figure 3, the cells infused with the synthetic RDS phytoalexin had an average amyloid plaque concentration of 14.837 pg/mL. Although this is greater than the amyloid plaque concentration of the cells infused with either natural phytoalexin extract, it is still significantly less than the positive control. This data sample also had the fewest outliers, indicating its reliability.

**Figure 3.**
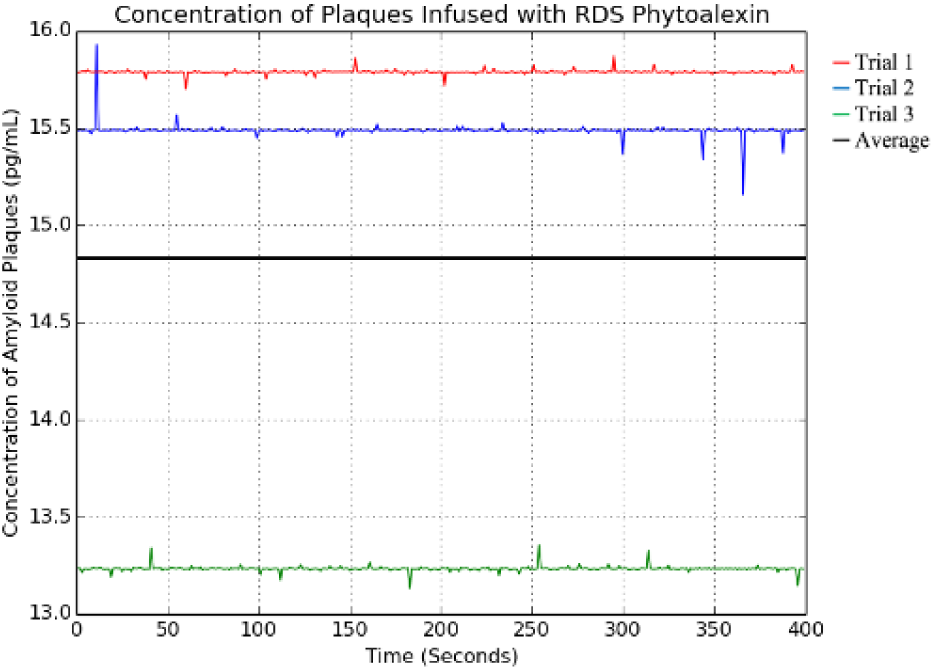
The concentration of amyloid plaques in the cells infused with the sythetic RDS phytoalexin extract over a time interval of 400 seconds.

### 2.3. Negative and Positive Controls

The negative control samples were not transfected with the ApoE4 plasmid, whereas the positive control samples were transfected with the plasmid; neither were in contact with phytoalexin treating agents.

The negative control samples, as shown in Figure 4a, contained neither the ApoE4 plasmid nor any phytoalexin treating agents. They only contained subcultured Neuro2a and therefore had the lowest measured amyloid plaque concentration, as there was no major factor to stimulate the accumulation of plaques. Due to the nature of spectroscopy, some of the readings taken were negative, as the spectrophotometer measured a lower absorbance compared to the baseline sample: water. This can be attributed to random error. Lastly, the positive control samples, as shown in Figure 4b, contained Neuro2a cells transfected with the ApoE4 plasmid. The samples did not contain any phytoalexin treating agents and thus had the highest concentration of amyloid plaques, an average of 34.073 pg/mL, as there were no treating agents to minimize the accumulation of the protein.

**Figure 4.**
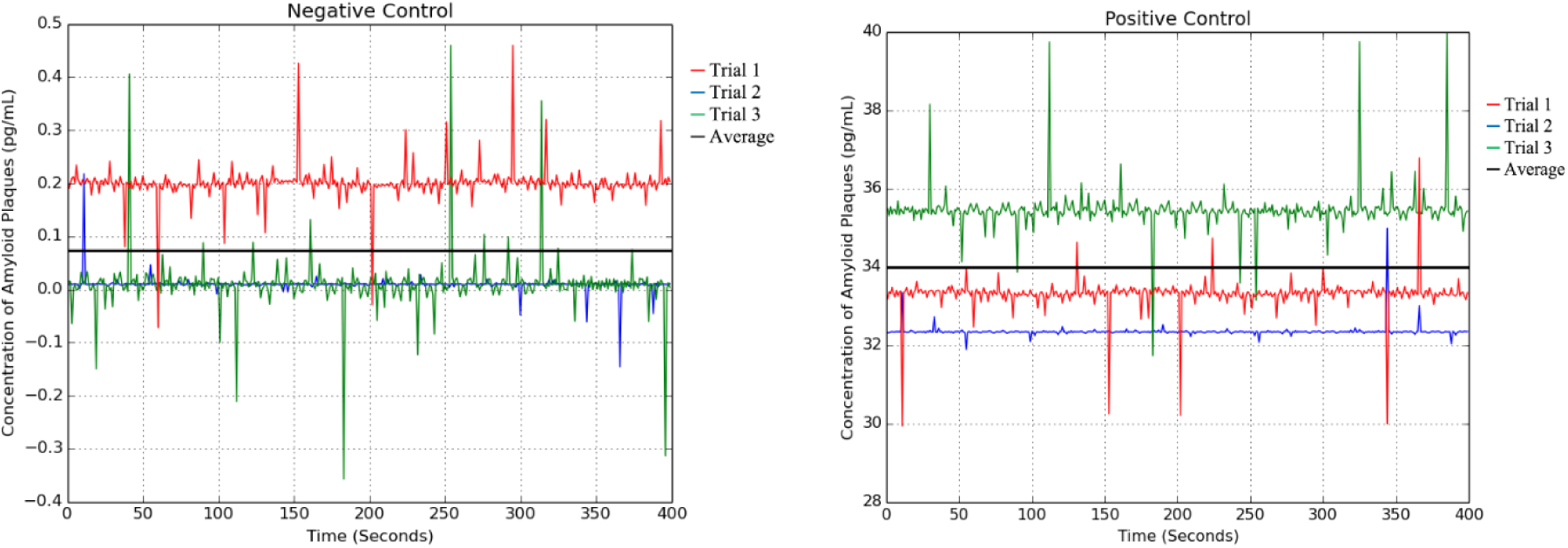
The concentration of amyloid plaques in the cells serving as the negative control (**a**) and the positive control (**b**) extract over a time interval of 400 seconds.

**Figure 5.**
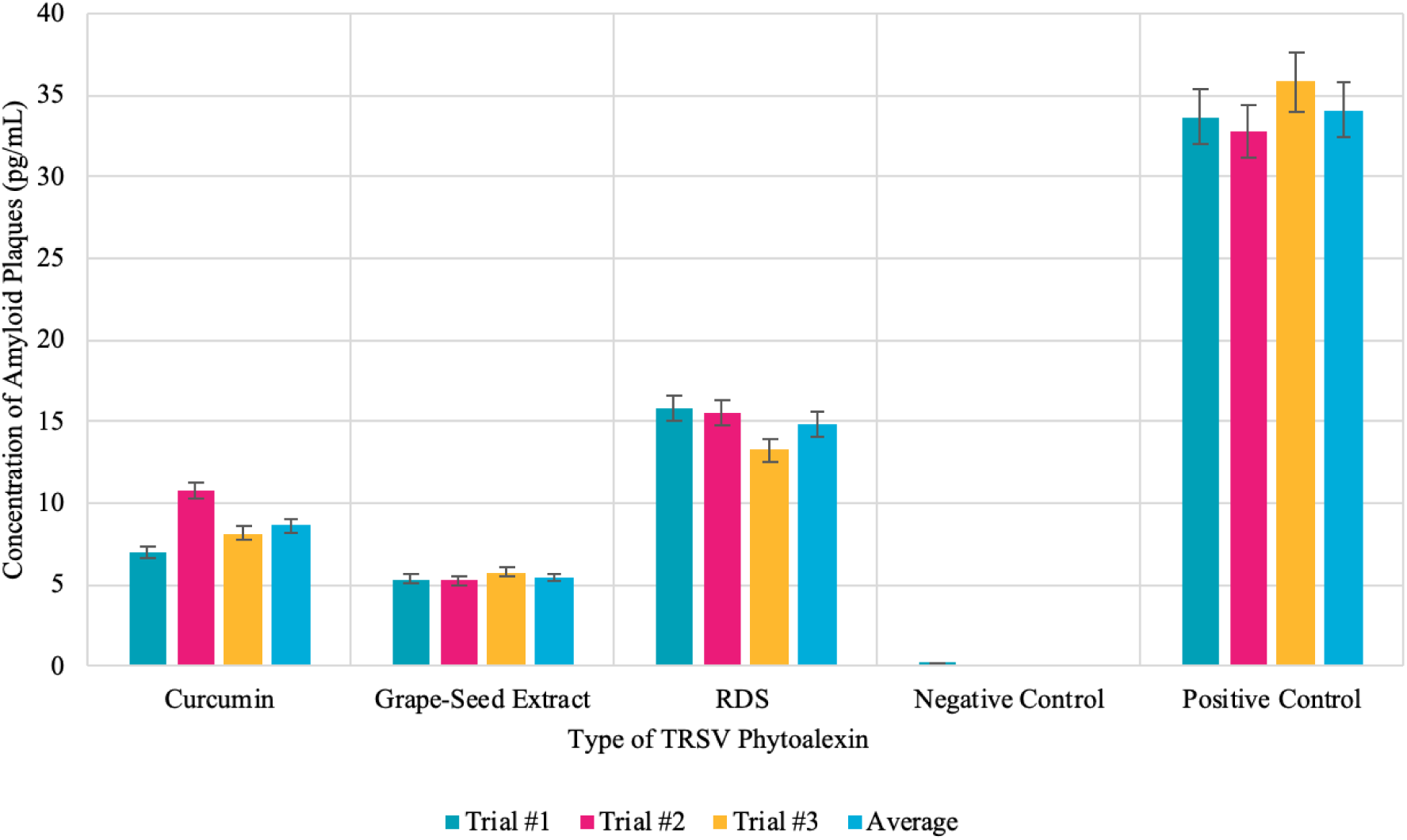
Graph of the concentration of amyloid plaques in the cells treated with both the natural and synthetic phytoalexin agents as well as both the positive and negative controls.

### 2.4. Summary Data and Analysis

Firstly, the negative control samples were not transfected with the ApoE4 plasmid, whereas the positive control samples were transfected with the plasmid; neither were in contact with phytoalexin treating agents. The negative control and the positive control samples had significantly different amyloid plaque concentrations: the negative with an average of 0.075pg/mL and the positive with an average of 34.073 pg/mL. This data is vital to this experiment, as it shows that the ApoE4 gene has a pivotal role in the accumulation of amyloid plaques within the Neuro2a cells. Secondly, grape seed extract counteracted the accumulation of amyloid plaques more effectively than curcumin extract. Additionally, both grape seed phytoalexin extract and curcumin phytoalexin extract were significantly more effective as treating agents compared to the synthetic RDS. It should also be noted that the error bars (set at standard error) of the samples do not overlap, indicating that the experimental results are significant. Lastly, standard deviation, standard error, and 95 percent confidence interval were calculated for each sample. All of these numbers are low, showing that the data is consistent and valid.

## 3. Materials and Methods

### 3.1. Preparing the Cells

To maximize cellular survival, the procedure was performed under optimal conditions (296K and seeding at 4×10,000 cells *cm*^2^) and in a sterile environment. A vial of Neuro2a (neuroblastoma) mouse cells was thawed using a water bath, and its contents were transferred to a 15mL centrifuge tube. 10mL of pre-warmed cell culture medium (Eagle Medium) was added to the centrifuge tube in a drop by drop manner to avoid osmotic shock. The cells were centrifuged at 200 x g for five minutes to allow for the removal of the freezing medium that contains dimethyl sulfoxide (DMSO).

The supernatant was discarded, and the pellet was re-suspended in 10mL of the complete, pre-warmed medium in a flask. 3mL of 100 units/mL penicillin, 100 *µ*g/mL streptomycin, 0.3 *µ*g/mL G418, 2.0 *µ*g/mL puromycin, and 1.0 *µ*g/mL tetracycline was each added to the flask to minimize the growth of unwanted bacteria. The flask was placed in the incubator with a rocking motion to ensure even distribution of the cells in solution. The cells were incubated at 37C in a humidified atmosphere containing 95 percent air and 5 percent CO2 until the cells have reached about 70-80 percent confluency (approximately 24 hours). This is when the cells are still growing exponentially (near the end of the log phase) and will result in improved overall cell viability, shorter lag time, and less aggregation.

### 3.2. Subculturing Adherent Cells

The culture was examined carefully for signs of contamination or deterioration and was handled with care during transport. The cells were rinsed using Dulbecco’s phosphate-buffered saline (DPBS), a balanced salt solution, by gently adding the solution (approximately 2 mL per 10 cm2 culture surface area) to the side of the flask opposite the attached cell layer to avoid disturbing the cell layer. This wash step removes any traces of serum, calcium, and magnesium that would inhibit the action of the dissociation reagent. The salt solution was removed using a pipette.

The pre-warmed dissociation reagent, TrypLE, was added to the side of the flask to remove cells from it. A microscope was used to confirm that the cells have released from the flask. When greater than 90 percent of the cells were detached, the vessel was tilted for a minimal length of time to allow the cells to drain. An equivalent of 2 volumes (twice the volume used for the dissociation reagent) of pre-warmed complete growth medium was added. The cells were transferred to a 15-mL conical tube and centrifuged at 200 x g for 10 minutes to remove any residual dissociation reagent. After centrifugation, the medium from the centrifuge tube was discarded with a pipette without disturbing the pellet. The pellet was resuspended with 10mL of warm, complete growth medium.

### 3.3. Calculating Viability Rates

After the hemocytometer was sterilized with 70 percent isopropyl alcohol, the coverslip was moistened with water affixed to the hemocytometer itself. The flask was gently stirred to ensure that the cells are evenly distributed. Before the cells have a chance to settle, 0.5 mL of cell suspension was taken out using a 5-mL serological pipette and placed into *µ*L of cells were placed into a new Eppendorf tube, and 400 *µ*L of Methylene Blue was added. After gently mixing, 100 *µ*L of Methylene Blue-treated cell suspension was taken out using a pipette and applied to the hemocytometer. Both chambers underneath the coverslip were filled very gently, allowing the cell suspension to be drawn out by capillary action.

Using a microscope, the gridlines of the hemocytometer were in focus with a 40X objective. The live, unstained cells were counted and recorded in one set of 16 squares. When counting, a system was employed whereby cells are only counted when they are set within a square or on the right-hand or bottom boundary line. Following the same guidelines, the dead cells stained with Methylene Blue were counted and recorded. Counting continued until all four sets of 16 corners are counted and recorded. Viability rates were calculated ((live cells/total cells)*100).

Using the percent viability of the cells, the cell suspension was diluted to the seeding density recommended for the cell line (4×10,000 cells *cm*^2^), and the appropriate volume of the medium into new cell culture flasks was pipetted. The cells were returned to the incubator. This procedure outlined in this subsection was repeated for each location on the flask (right-bottom, center-bottom, left-bottom, right-top, center-top, left-top, center-center) three times to ensure accuracy of cellular viability throughout different regions of the flask. After 24 hours in incubation, the cells were evenly divided into five 20mL different flasks. Each of the flasks was labeled as the positive control, negative control, grape seed extract test, curcumin extract test, and RDS (resveratrol dietary supplement) test.

### 3.4. APOE-4 Plasmid DNA Transfection

The cells were transferred from the five flasks to five 3-well plates and were distributed evenly throughout all of the wells. After labeling the well-plates respectively based on the treating agent or control type, 24 *µ*L of Opti-MEM medium and 4*µ*L of Lipofectamine LTX was added to each tube using an 8mm insulin syringe. Five other Eppendorf tubes were prepared with 250*µ*L of Opti-MEM medium each. 5*µ*L of APOE-4 Bacterial Plasmid DNA was added to each of the tubes (DNA concentration is at 1*µ*g/*µ*L). 5*µ*L of Plus Reagent was also added to each of the four tubes. These five tubes *µ*L from each of the five diluted plasmid DNA tubes were added to each of the 24 Eppendorf tubes that were prepared earlier in this section. These tubes serve as the DNA-reagent complex. The cells in the well plates were incubated at 37C for five minutes. After incubation, 50*µ*L of the DNA-reagent complex was added into all of the wells of four out of the five wells. The DNA-reagent complex was not added to the well-plate labeled as “negative control.” Place the well-plates back into the incubator at 37C for 48 hours.

### 3.5. Resveratrol Treatment and Bradford’s Assay

After incubation, all five 3-well plates were removed from the incubator and were placed in the cell culture hood. For the well plate labeled as ‘grape seed extract test,’ 30*µ*L of grape seed extract was added to each of the three wells in the well plate. For the well plate labeled as ‘curcumin extract test’, 30*µ*L of curcumin extract was added to each of the three wells in the well plate. For the well plate labeled as ‘RDS test,’ 30*µ*L of RDS was added to each of the three wells in the well plate. The well-plates that are labeled as ‘positive control’ and ‘negative control’ were not treated. The well-plates were placed into the incubator at 37C for 1 hour.

To determine beta-amyloid protein concentration via Bradford’s assay, Bradford’s reagent was first prepared by dissolving 100 mg of Coomassie Brilliant Blue G-250 in 50 ml of 95 percent ethanol. This was filtered through Whatman 1 paper just before use. 5*µ*L of Bradford’s dye reagent was added to each of the six wells in each well plate and were incubated for 5 min at 37C. 5mL of the solution was transferred from each of the three wells into three different cuvettes. This was done for all five well-plates. An Ultraviolet-visible spectrophotometer measured the absorbance at 595 nm for 400 seconds of the different samples in the cuvettes. The readings were run through a low-pass filter in Python to generate cleaner data with statistical analysis. Cultures were disposed of using proper bleach methods. Each of the wells from the well plates represents a trial for a total of three trials for each of the three agents and the two controls.

## 4. Conclusions

The experimental data showed, based on the amyloid plaque concentration measured, that the grape seed extract was the most effective phytoalexin treating agent compared to the curcumin extract and RDS. RDS was the least effective treating agent. It is also essential to identify that all of the tested phytoalexins, regardless of natural or chemical composition, lessened the accumulation of amyloid plaques to some degree compared to the positive control.

Experimental errors are possible, primarily due to the nature of the experiment. One possible error is the presence of microscopic particles on the sides of the cuvettes that can alter the absorbance measured by the spectrophotometer and ultimately return an inaccurate value for the concentration of the amyloid plaques. This error was minimized as much as possible by using non-static wipes to clean the cuvette. Another potential error is that ApoE4 plasmid did not properly assimilate into the Neuro2a cells because of contaminants that might hinder the plasmid transfection and gene expression. Although improbable, this would result in the creation of an inaccurate model for that specific sample. By working in a sterile laminar hood, this was minimized as much as possible. However, this error could be minimized further by conducting the experiment in a positive pressure room.

This project has great practical relevance: it not only evaluates the overall treatment capabilities of phytoalexins, but it also identifies which natural and synthetic TRSV phytoalexin specifically serves as the best limiter of amyloid plaque accumulation, the cause of many of the neurodegenerative effects commonly associated with AD. Additionally, this project determines how a synthetic phytoalexin compares to natural phytoalexins with regard to treatment ability. Because the accumulation of amyloid plaques plays a central role in the pathogenesis of AD, this experiment also sheds light on potential treatments that can be further explored in future in-vitro or clinical experimentations. Lastly, this methodology defines a new model of studying the treatment capabilities of substances and compounds that may be used in future AD treatment.

## Acknowledgments

I would like express my deep gratitude and sincere thanks to my mentors and instructors, Mrs. J. Naughton, Mrs. J. Baylor, and Mrs. L. Turngren, for their valuable guidance and advice.

## Abbreviations

The following abbreviations are used in this manuscript:

AD: Alzheimer’s Disease
DPBS: Dulbecco’s Phosphate-Buffered Saline
DMSO: Dimethyl Sulfoxide
CBB: Coomassie Brilliant Blue
TRSV: Trans-Resveratrol
RDS: Resveratrol Dietary Supplement
APOE: Apolipoprotein E
Aβ: Beta Amyloid
APP: Amyloid Precursor Protein
ROS: Reactive Oxygen Species
AICD: APP Intracellular Domain

